# Ageing blunts the phospho-proteomic response to resistance exercise in humans

**DOI:** 10.64898/2026.06.02.729449

**Authors:** Andrew S. Palmer, Isabelle Alldritt, William Apro, Hannah Huckstep, Oscar Horwath, Lina H. H. Le, Tabitha Cree, Sarah E. Deemer, Emma L. Watson, Richard J. Mills, Sean J. Humphrey, Andrew Philp

**Author notes:** **Corresponding author** Correspondence to Andrew Philp. These authors contributed equally: Andrew S. Palmer, Isabelle Alldritt, William Apro.

## Abstract

Ageing blunts the adaptive growth response in human skeletal muscle. To define the molecular processes underlying this impairment, we performed unbiased analysis of the phosphoproteome and total proteome in healthy young and old human skeletal muscle following acute resistance exercise (ResEX) and essential amino acid (EAA) ingestion. Ageing led to global suppression of the growth-induced global phosphoproteome, despite intact mTORC1 activation in old muscle. Our results identify widespread effects of ageing on skeletal muscle growth and highlight novel pathways for therapeutic development in aged skeletal muscle.

## Main

Skeletal muscle deterioration is a common hallmark of ageing, with muscle strength and size known to decrease progressively from midlife^1^. When sustained, this loss leads to the onset of sarcopenia^2^ and can progress to loss of functional independence and frailty^3^. It has been proposed that a reduced sensitivity to growth stimuli, such as resistance exercise and protein ingestion, underlie sarcopenia progression, in a process termed anabolic resistance^4,5^. However, whether ageing per se, or other lifestyle factors such as physical inactivity or poor dietary practice act as the primary driver of anabolic resistance is still under debate^6^. We recently reported that when ResEX and EAA intake are optimally prescribed, muscle protein synthesis rates and mTORC1 signalling in old healthy men increase in a response comparable to young counterparts^7^. Therefore, this model allows us to directly test the role of ageing on the molecular responses to an optimal growth response.

We performed unbiased liquid chromatography tandem mass spectrometry (LC-MS/MS) analysis of the skeletal muscle phospho/total proteome in 10 old (70 ± 3yrs) and 10 young (22 ± 3yrs) male participants at rest and following acute unilateral resistance exercise (ResEX) with EAA ingestion as previously described (Fig. 1A)^7^. Young males (23.4 +-0.8 kg/m2) and old males had comparable body mass index but young males (58 +-6kg) had greater leg strength than old males (30 +-7kg) (Fig. 1A)^7^. After preprocessing and filtering, we quantified 17,518 phosphosites (located on 3,561 unique proteins) and 4,444 proteins across all samples. Using principal component analysis (PCA), we observed that both the phosphoproteome and proteome separated by age along the first principal component with 8.4% and 13.7% of the variation explained along component 1 (Fig. 1B).

**Figure 1.**
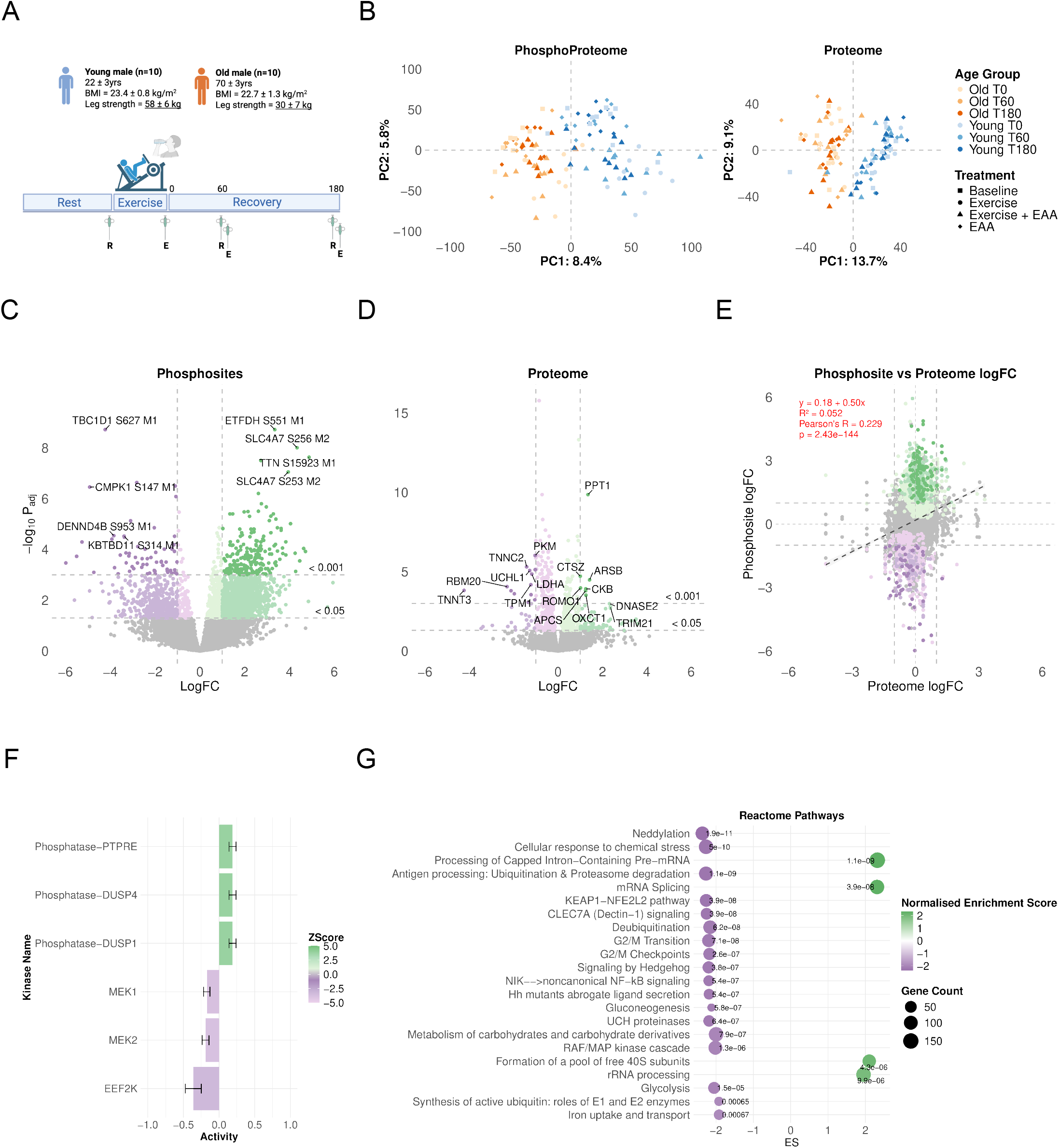
Old vs Young analysis. **A.** Study overview. **B**. Principal component analysis of both the phosphoproteome and proteome. Old participants and timepoints indicated by orange points and young participants by blue points. Square points represent baseline samples, circle points exercise only, triangle points exercise with essential amino acid (EAA) supplementation, and diamond points only EAA supplementation. **C-D**. Identified differentially regulated phosphosites and proteins from an Old vs Young comparison at baseline. X-axis is the log fold change and Y-axis −log10 adjusted p-values. **E**. Phosphosite vs proteome logFC plot with proteome logFC on the Y-axis and phosphosite logFC on the X-axis. Grey dashed lines on all plots indicate logFC |1| or adjusted P value cut off <0.05 and <0.001, and all purple points indicate downregulated and green upregulated **F**. Inferred kinase and phosphatase activity Z-scores estimated using RoKAI based on phosphoproteomic measurements between Old and Young at baseline. The heatmap displays Z-scored kinase/phosphatase activity, where positive Z-scores (green) indicate increased activity and negative Z-scores (purple) indicate reduced activity, activity measure is on the X-axis and the kinases or phosphatases are those identified with FDR < 0.05. **G**. The top Reactome pathways identified using functional gene set enrichment analysis using the protein logFC from the Old vs Young comparison. X-axis indicates the normalised enrichment score and gene counts are indicated by point size. FDR p values are displayed as text. Down regulated and up regulated pathways indicated by purple and green respectively.

To determine baseline differences between old and young muscle protein signalling, we performed differential analysis of the phospho and total proteome from the resting biopsy. We observed 1,569 upregulated and 466 downregulated phosphosites in old vs young muscle with FDR < 0.05 and logFC > 1 (Fig. 1C). Only 61 proteins were upregulated and 32 proteins downregulated with FDR < 0.05 and logFC > 1 (Fig. 1D). When proteins with FDR < 0.05 and any logFC were included, there were 416 upregulated and 277 downregulated proteins. We observed a weak positive correlation between the differential abundance of phosphosites and their corresponding total proteins in old versus young muscle (R=0.23, R^2^=0.052, P < 0.001), indicating that only 5.2% of the variance in phosphorylation changes is explained by changes in protein abundance (Fig. 1E). We performed kinase activity assessments on the phosphosites using RoKAI^8^, finding increased activity of the phosphatases PTPRE, DUSP1 and DUSP4, and decreased activity of MEK1, MEK2 and EEF2K with age (Fig. 1F). Gene set enrichment analysis of the proteins revealed decreased enrichment of Reactome pathways involved in energy metabolism, ubiquitin-proteasome system, growth and regeneration and an increased enrichment in mRNA and rRNA processing pathways (Fig. 1G). Similarly, we observed differences in phosphorylation patterns between old and young muscle, with changes in kinase activity and altered enrichment of energy metabolism, growth, and regeneration pathways indicating sustained alterations in the homeostasis of resting muscle with age.

Next, we quantified the global phosphoproteomic response to ResEX and EAA intake to determine age-related differences in anabolic signalling. Young muscle demonstrated robust phosphoproteome signalling, with ResEX alone triggering 2,414 upregulated and 881 downregulated phosphosites immediately post exercise (FDR <0.05) (Fig. 2A). When combined with EAA intake, young muscle maintained an increased signalling at both 60 and 180mins (1,817 up; 802 down at T60; 1,620 up; 327 down at T180) (Fig. 2A). EAA supplementation alone produced more modest signalling at 60 and 180mins (438 up, 184 down at T60; 374 up, 180 down at T180) compared to ResEX+EAA (Fig. 2A). In contrast, old muscle displayed markedly lower responses across the same conditions: ResEX alone induced approximately half as many upregulated phosphosites as the young muscle (1,210 up, 798 down); and this persisted with ResEX+EAA at 60 and 180mins (1,182 up, 563 down; 550 up, 577 down at T180)(Fig. 2B). EAA alone produced similar signalling in old and young at 60mins but was lower in old muscle at 180mins (367 up, 262 down, 163 up, 244 down)(Fig. 2B). To investigate age-dependent responses in phosphoproteome signalling, we analysed interaction effects to identify age-specific phosphorylation. The number of age-specific phosphosites in old versus young muscle (FDR <0.05) were: ResEX at T0 (232 up, 508 down), ResEX+EAA at T60 (180 up, 234 down), and ResEX+EAA at T180 (139 up, 592 down)(Extended Data Fig. 1A), indicating more downregulated and dampened phosphorylation in old muscle compared to young muscle. These findings indicate a pronounced age-related decline in global phosphoproteomic responsiveness, with old muscle exhibiting both a reduced magnitude and altered temporal profile of signalling across ResEx and EAA conditions(Fig. 2C).

**Figure 2.**
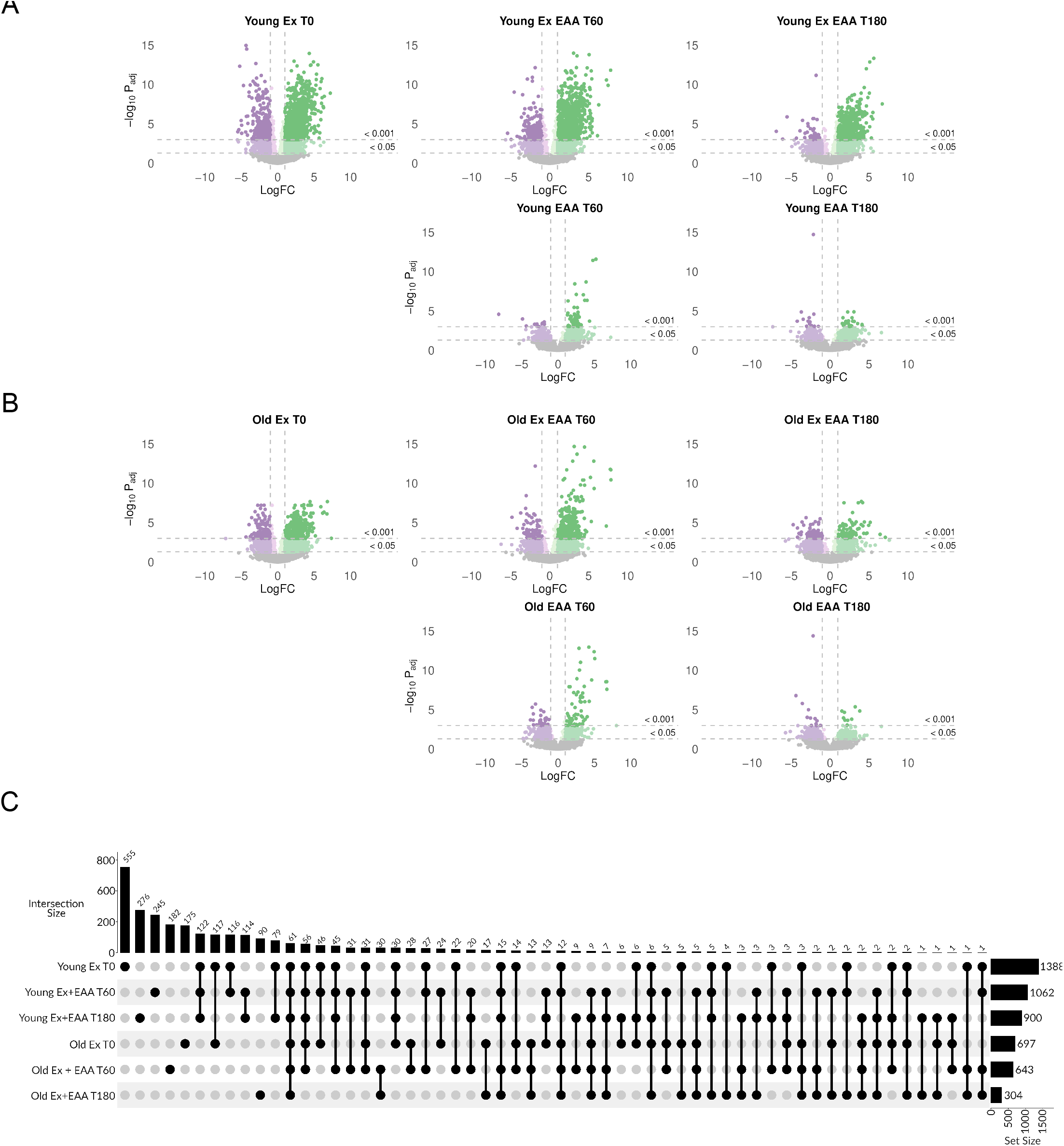
Exercised conditions vs baseline in young and old participants. **A.** Identified phosphosites from Timepoint (T)0, T60 and T180 with either exercise (Ex) or both exercise and essential amino acid (EAA) supplementation compared with baseline in young participants. X-axis is the log fold change and Y-axis −log10 adjusted p-values. **B**. Identified proteins from T0, T60 and T180 with either Ex or both Ex and EAA supplementation compared with baseline in old participants. X-axis is the log fold change and Y-axis −log10 adjusted p-values. **C** UpSet plot showing all differentially regulated phosphosites with adjusted P values < 0.05 and absolute log fold change > 1.5.

A parallel analysis of the global proteome revealed minimal changes with ResEX or EAA intake, with only 11 proteins (6 upregulated, 5 downregulated) altered across all comparisons in both young and old muscle (Extended Data Fig. 2A–B) and no proteins identified in interaction models (Extended Data Fig.1B). We compared the logFC of phosphosites with those of their corresponding proteins and found no meaningful correlation (R = 0.02–0.09, R^2^ = 0.002–0.009), indicating that the observed phosphorylation changes are largely independent of changes in protein abundance (Extended Data Fig. 1C, Extended Data Fig. 3A-B).

To understand the hierarchical regulation of the age-dependent phosphoproteome, we assessed kinase activity using RoKAI^8^. We observed four main clusters of kinases with distinct activity patterns across all comparisons (Fig. 3A). Two clusters displayed similar kinase activity, and two clusters were divergent in kinase activity with age. Despite an overall suppression of the global phosphoproteome in aged muscles, specific kinases such as p70S6K, p90RSK, mTOR, and AKT2 maintained similar activity levels in both young and old individuals after ResEX and EAA ingestion (FDR < 0.05, Fig. 3A). Similar phosphorylation patterns on the mTOR pathway were observed in both young and old muscles (Extended Data Fig. 4A), aligning with prior results that indicated sustained mTORC1 activity and muscle protein synthesis rates in healthy older men after optimal exercise and EAA ingestion^7^. In contrast, aged muscles showed reduced activity in JNK1-3, p38A, p38D, p38G, PAK1-2, CDK1, and CDK5 (only young muscles exhibited increased activity, FDR < 0.05) after ResEX and EAA intake. Conversely, there was an increase in MEK1, MEK2, MKK6, ERK2, and PKM iso2 activity in old muscles post ResEX and EAA intake (only old muscles showed increased activity, FDR < 0.05). This suggests dysregulation in protein signalling pathways with age. Additionally, we observed decreased activity with age in the phosphatases PTPRE, DUSP1, DUSP4, PTPN1, PPP2CA, PPP2CB and PTPN7 in response to exercise (Fig. 3B) in old muscle (FDR < 0.05). Differing phosphorylation patterns on the MAPK pathway were observed in old compared with young muscles (Extended Data Fig. 4B).

**Figure 3.**
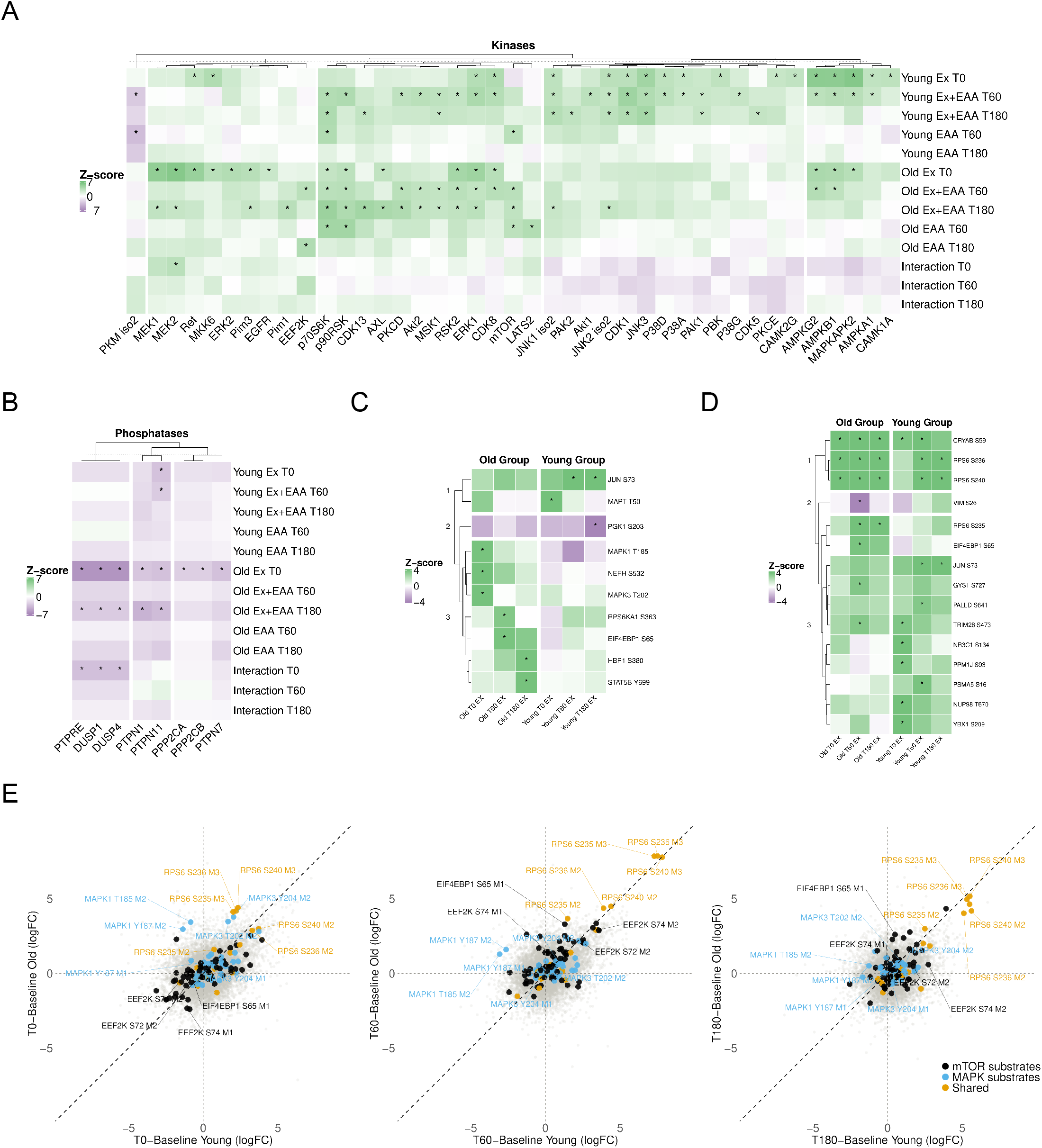
Regulated phosphosite analysis. **A-B.** Heatmap of inferred kinase and phosphatase activity Z-scores across experimental conditions. Activity was estimated using RoKAI based on phosphoproteomic measurements. The heatmap displays Z-scored kinase/phosphatase activity, where positive Z-scores (green) indicate increased activity and negative Z-scores (purple) indicate reduced activity. Kinases/phosphatases were hierarchically clustered based on Euclidean distance. Asterisks indicate kinases or phosphatases with FDR < 0.05. **C**. Heatmap of substrate Z-scores across experimental conditions for cluster 1 kinases. The heatmap displays Z-scored phosphosite fold changes, hierarchically clustered based on Euclidean distance. Asterisks indicate phosphosites with FDR < 0.05. **D**.Heatmap of substrate Z-scores across experimental conditions for cluster 3 kinases. The heatmap displays Z-scored phosphosite fold changes, hierarchically clustered based on Euclidean distance. Asterisks indicate phosphosites with FDR < 0.05. **E**. Young phosphosite logFC vs. old phosphosite logFC for *T*_0_, *T*_60_, and *T*_180_. The black dashed line is the unit diagonal (*y* = *x*), representing equal fold-change between age groups. Points are color-coded by kinase substrate specificity: black dots represent mTOR substrates, blue dots represent MAPK substrates, and yellow dots represent substrates shared by both kinases.

We characterised the age-related phosphoproteomic response to ResEx and EAA to identify the key substrates driving altered kinase activity. We found that cluster one (Fig. 3A) was defined by an increased phosphorylation cascade of MAPK1 (T185), MAPK3 (T202), RPSKKA1 (S363), HBP1 (S380), and STAT5B (Y699) (Fig. 3C) in the old group. Whereas, cluster three consisted of similar growth signalling across age groups on RPS6, but blunted stress response signalling on JUN (S73) and NR3C1 (S134) with age (Fig. 3D). Finally, to visualise the age-dependent phospho response, we generated logFC vs logFC plots for each timepoint. RPS6 serines 235, 236 and 240 had greater phosphorylation at the acute time point in the old group, before increasing to similar levels at T60 and T180 after Ex and EAA intake in both groups (Fig. 3E). MAPK1 phosphorylation on threonine 185 and tyrosine 187 increased in the old group however, decreased in the young at the acute and T60 timepoints. Collectively, these findings indicate that while the mTOR pathway remains active in response to ResEX and EAA stimuli during ageing, the activities of the JNK/P38 and MAPK/ERK pathways are altered with age.

In summary, our study reveals that when healthy young and old male individuals perform a matched bout of ResEX with EAA ingestion, there is a blunted phosphoproteomic response in aged muscle, independent of changes in mTORC1 signalling or changes at the level of total proteome. Specifically, ageing led to alterations in MAPK signalling in parallel with reduced activity of numerous phosphatases. Collectively, these findings indicate that while mTORC1 signalling remains sensitive to ResEX and EAA during ageing, the activities of JNK/P38 and MAPK/ERK pathways are altered with age, suggesting that despite similar growth responses there may be an altered stress remodelling response with age. Our unbiased approach allowed us to determine the global phosphoproteomic response to ResEX+EAA, uncovering novel pathways that may underlie the loss of skeletal muscle size and strength. However, the study design does not allow us to pinpoint how ageing alters these pathways, nor the contribution of each to the adaptive growth response. Addressing these questions may uncover novel approaches to promote muscle growth in aged skeletal muscle.

## Methods

### Ethical Approval

Participants were recruited at the University of Birmingham and the Swedish School of Sport and Health Sciences. The study was approved by the West Midlands, Black Country Research Ethics Committee, #17/WM/0068 and the Swedish Ethical Review Authority #2017/2107-31/2 and performed in accordance with the Declaration of Helsinki. All participants provided written informed consent before participation.

### Participants and Study Design

The samples used in this study have been described previously^7^. Briefly, ten young (18-35) and ten old (65-75), non-smoking, men were recruited for this study, and were excluded if they had cardiovascular or metabolic disease or regularly participating in structured resistance exercise. On the day of the experiment, participants arrived at ∼7:00AM and received a constant infusion of L-[ring-^13^C6]-phenylalanine for the entire experimental period. After 150min resting muscle biopsies were collected from the *m. vastus lateralis* using the Bergström needle technique. Participants performed unilateral resistance exercise consisting of 3 warm-up sets with 2-min rest intervals followed by 10 sets at their predetermined 10RM with 3-min rest intervals. Immediately after exercise, a biopsy was collected from the exercise leg, after which participants consumed an essential amino acid drink. During recovery biopsies were obtained from both the exercised and resting leg at 60mins and 180mins after consumption of the essential amino acid drink. Biopsies were immediately frozen in liquid nitrogen and stored at −80°C.

### LC-MS/MS phosphoproteomes

#### Sample preparation

Frozen, powdered muscle biopsies (20 mg wet weight) were lysed in 4% SDC buffer (4%Sodium Deoxycholate/100 mM Tris pH 8.5) with boiling at 95°C for 5 min (1,500 rpm), followed by sonication (tip-probe sonicator, 30% output, 3s on/3s off, 15 s total). Samples were centrifuged at 14,000 xg for 15 min at room temperature, and supernatant collected into clean tubes. A 5 µL aliquot of protein was taken and diluted with 45 µL 8M Urea for protein content estimation by BCA assay, and 50 µg of protein in 180 µL was used for phosphoproteome and proteome analysis. Protein was reduced and alkylated by the addition of 20 µL Tris (2-carboxyethyl)phosphine (TCEP) and 2-chloroacetamide (CAA) (10 mM and 40 mM final concentration, respectively) with heating at 45 °C for 5 min, after which samples were incubated at room temperature for 40 minutes. Lysates were precipitated by the addition of 800 µL (four volumes) of methanol and 200 µL (one volume) of chloroform, with mixing in-between, followed by the addition of 600 µL (three volumes) of water and further mixing. Samples were centrifuged at 3,000 xg for 5 minutes at room temperature, after which the top 80% of the supernatant (upper phase) was carefully removed. To the remaining liquid, 600 µL methanol was added with brief mixing, and samples were centrifuged at 3,000 xg for 5 min at room temperature. Supernatants were removed and discarded, and protein pellets were briefly air-dried with inversion. Protein was resuspended in 4% SDC buffer with sonication and NaCl added to 10 mM. Protein was subsequently precipitated by the addition of 4.2 volumes of −30°C acetone with mixing, and samples placed at −30°C overnight. Samples were centrifuged at 20,000 xg for 15 min at 4°C, supernatants decanted and discarded, and pellets briefly air-dried with inversion. Protein pellets were resuspended in 155 µL of 4% SDC buffer with mixing (5 min, 2,000 rpm), followed by sonication (30% output, 3s on/3s off, 15s total). A 5 µL aliquot was taken and diluted with 45 µL 8M Urea for protein estimation by BCA assay. After determining protein yield, all samples were diluted to equal concentration in a final volume of 150 µL of 4% SDC lysis buffer and stored for further processing. Protein samples containing 50 µg protein in 150 µL lysis buffer were subsequently digested by the addition of 1:50 Trypsin + LysC overnight (18 hr, 37°C, 1,500 rpm), and phosphopeptides enriched using the EasyPhos method, as previously described^9^. Peptide ‘flow-through’ from phosphopeptide enrichments were retained and used for proteome analysis.

### LC-MS/MS measurement

Enriched phosphopeptides in MS loading buffer (2% ACN, 0.3% TFA) were loaded onto in-house fabricated 55-cm columns of 75-µm I.D. packed with 1.9-µm ReproSil-Pur C18-AQ particles, using a Vanquish Neo UHPLC coupled to an Orbitrap Astral mass spectrometer (Thermo Fisher Scientific). Column temperature was maintained at 60 °C in a Sonation column oven. Peptides were separated using a binary buffer system of 0.1% formic acid (buffer A) and 80% ACN, 0.1% formic acid (buffer B) at 400 nl min^−1^, with a gradient of 3– 19% B over 20 min and 19–41% B over a further 10 min (30-min effective gradient), followed by a wash at 99% B from 31–39 min and re-equilibration, for a total cycle of 40 min. Ionisation was achieved by nanoelectrospray (NSI) at 2.4 kV. Peptides were analysed with one full MS scan in the Orbitrap (380–980 m/z, R = 240,000 at m/z 200, normalised AGC target 500% / 5 × 10^6^, maximum injection time 5 ms), followed by 300 data-independent acquisition (DIA) MS/MS scans (380–980 m/z, 2 m/z isolation windows) with HCD fragmentation (NCE 24%) and fragments were recorded in the Astral analyser (150– 2,000 m/z, target 5 × 10^4^, maximum injection time 3.5 ms).Enriched phosphopeptides in MS loading buffer (2% ACN, 0.3% TFA) were loaded onto in-house fabricated, 50-cm columns with a 75-µm I.D. and packed with 1.9 µM C18 ReproSil Pur AQ partcles using a Dionex U3000 HPLC coupled to an Orbitrap Exploris 480 mass spectrometer (Thermo Fisher Scientific). Column temperature was maintained at 60 °C in a Sonation column oven, and peptides separated using a binary buffer system comprising 0.1% formic acid (buffer A) and 80% ACN plus 0.1% formic (buffer B), at a flow rate of 400 nl nmin–1 with a gradient of 3–19% buffer B over 80 min followed by 19–41% buffer B over 40 0 min, resulting in ∼2-h gradients. Peptides were analysed with one full scan (35030–1,400 m/z, R = 120,000) at a target of 3 × 106 ions, followed by 48 data-independent acquisition (DIA) MS/MS scans (3503–1,022 m/z) with HCD (target 3 × 10^6^ ions, maximum injection time 22 ms, isolation window 14 m/z, 1 m/z window overlap, NCE 252%), with fragments detected in the Orbitrap (R = 15,000).

### LC-MS/MS total proteomes

#### Sample preparation

Approximately 8 µL of ‘flow-through’ material containing 1 µg of peptides was retained from phosphopeptide enrichment for total proteome analysis. To desalt peptides, two stacked Empore SDB-RPS disks (Sigma-Aldrich, USA) were added to a 200 µL pipette tip as described^10^. To the 8 µL peptide samples, 92 µL of water was added, followed by 100 µL 99% ethyl acetate/1% TFA, and mixed thoroughly. Samples were loaded on SDB-RPS StageTips and spun through to dryness. StageTips were washed with 100 µL 99% isopropanol/1% TFA, then once with 100 µL 0.2% TFA/5% acetonitrile. Peptides were eluted into a 96 well plate using 5% ammonium hydroxide/25% acetonitrile before being dried under vacuum at 45°C (Eppendorf Concentrator Plus, Eppendorf, Germany). Peptides were reconstituted in 5% formic acid and stored at 4°C until analysis.

### LC-MS/MS measurement

Proteomics analysis was performed on a Vanquish Neo 1 UHPLC pump and autosampler coupled to a Q-Exactive HF-X mass spectrometer (Thermo Fisher Scientific, USA). 2µl sample was injected for a total concentration of 5ng. Peptides were resolved over a gradient from 5% to 98% 0.1% formic acid in 80% ACN over 75 minutes with a flow rate of 0.3µl/min. MS1 scans were acquired from 350 to 1650 m/z (60,000 resolution, 3e6 AGC target and 50ms maximum injection time). MS2 scans were acquired using a data independent acquisition method. Measurements of bovine serum albumin (BSA) were taken at the start and end of the run for quality control.

#### Phosphoproteome preprocessing

The following preprocessing and analyses were performed in the R software environment version 4.5.1 (v.4.5.1)^11^. To remove unreliable phosphosites, all phosphosites that had a localisation probability of < 0.75 or were missing in at least 50% of one condition were removed and all values < 3 were converted to missing values as they were considered unreliable measurements indistinguishable from background noise. Raw intensities were log2 transformed, after which we performed centre-median normalisation. For conditions with at least 2 samples with measured values, we imputed missing values by group sampling from a down shifted normal distribution (down shifted mean = group mean – 1.6x replicate standard deviation with a down shifted standard deviation = 0.6 x group standard deviation) and set a minimum threshold of log2(3.0) to match our measurement cut off during filtering. The remaining missing values were imputed by a global down shifted normal distribution (with the same down shifted and minimum threshold properties).

#### Proteome preprocessing

Raw data files were processed using Spectronaut^12^ (version 19.6, Biognosys AG), with “.directDIA^TM^”. A spectral library was generated against the human UniProt FASTA database (accessed in January 2025 containing 83,607 entries and 20,659 genes) with the inclusion of common contaminants and iRT standards for protein identification. Default settings were used for search parameters. The false discovery rate (FDR) for peptide-spectrum matches and protein identifications was set to 1% at peptide level. Data analysis was performed in R v4.5.1. Proteins identified as contaminants, decoy sequences, or detected in fewer than three replicates were excluded from further analysis. Label-free quantification (LFQ) intensities were log2-transformed and normalised using a median-centreing approach. Missing values were imputed assuming a missing-not-at-random pattern, with imputation performed by drawing random values from a left-shifted Gaussian distribution (1.8 standard deviations, width 0.3).

#### PCA analysis

Principal component analysis was used to determine major sources of variation in the phosphoproteome and proteome data using the prcomp function in R v4.5.1^11^.

#### Differential analysis

Differentially regulated phosphosites and proteins were identified using the limma package v3.64.3^13,14^ by calculating empirical bayes moderated t-statistics. Duplicate correlations were included in the model to account for the paired experimental design. Main effects of each condition were calculated as the difference to the resting baseline sample. Interaction effects (age x intervention) were calculated for T0, T60, and T180 to assess if the response to the intervention (ResEX or EAA) was different between young and old. To correct for multiple hypothesis testing, we used the Benjamini and Hochberg (BH) False Discovery Rate (FDR) method^15^ with a cutoff of <0.05.

### Kinase activity analysis

ROKAI^8^ was used to infer kinase activity using functional networks. Data was prepared for uploading in a locally running ROKAI app v2.2.0 using shiny v1.11.1. The following settings were applied: Reference Proteome = Uniprot Human, Fold Changes = Centered, Kinase Substrate Dataset = PSP+Signor, RoKAI Network = KS+PPI+SD+CoEv, Phosphatase Substrate Dataset = DEPOD. We used sites in the functional neighbourhood and included phosphatases in the analysis. For reporting and visualisations we kept kinases and phosphatases with a minimum of 3 substrates. To determine phosphosites underlying the kinase activities, we extracted substrate ZScores and FDR values from the kinase targets tables generated by ROKAI.

### Functional gene set enrichment

The FGSEA package^16^, v1.34.2, was used to conduct functional gene set enrichment of differentially regulated proteins in Reactome gene sets, excluding those gene sets with <5 or >500 genes. We used all proteins in the differential protein analysis as a background list. The collapsePathways function from FGSEA was then applied to reduce redundancy in the enriched pathways list.

### Data visualisations

PTMNavigator^17^ was used to overlay phospho and total proteome data with the mTOR (HSA04150) and P38 MAPK (WP400) pathway diagrams. The following R packages were utilised for the remaining data visualisation generation, ggplot2 v3.5.2, ggrepel v0.9.6^18^, ComplexHeatmap v2.24.1^19^, cowplot v1.2.0^20^ and multipanelfigure v2.1.6^21^.

## Data availability

Phosphoproteomic and proteomic data have been deposited to the ProteomeXchange http://proteomecentral.proteomexchange.org Consortium via the PRIDE^22^ partner repository with the dataset identifiers PXD078824 (phosphoproteome) and PXD074415 (proteome). All reported phosphosites, proteins, kinases, phosphatases, rokai kinase tables and targets and reactome sets are deposited in the Zenodo repository^23^ https://doi.org/10.5281/zenodo.20387916.

## Code availability

All code used for this publication and a list of all R packages loaded during data analyses including version numbers are available in the Zenodo repository^23^https://doi.org/10.5281/zenodo.20387916.

## Acknowledgements

We thank Professor Mirana Ramialison, Professor Sharon L. Naismith, Professor Susan E. Kurrle and Dr Mark Larance for their help and support throughout the course of the study. This project was supported by funding from the European Research Council (ERC) under the European Union’s Horizon 2020 research and innovation programme (grant agreement no. 707336) to A.P. The Åke Wiberg Foundation (grant no. M17-0259) and The Lars Hiertas Memorial Foundation (grant no. FO2017-0325) to W.A. The Elisabeth and Gunnar Liljedahl Foundation to O.H. and by Wellcome Leap’s Dynamic Resilience Program (jointly funded by Temasek Trust).

R.J.M. is supported by National Health and Medical Research Council (NHMRC) grants (GNT2013189 & GNT2049456), and is a member of The Novo Nordisk Foundation Center for Stem Cell Medicine, reNEW, which is supported by Novo Nordisk Foundation grant no. NNF21CC0073729. S.J.H. is supported by a National Health and Medical Research Council (NHMRC) Investigator Grant (2026905), Synergy Grant (2035975) and is a member of The Novo Nordisk Foundation Center for Stem Cell Medicine, reNEW, which is supported by Novo Nordisk Foundation grant no. NNF21CC0073729.

## Contributions

Conceptualisation: W.A., A.P.; Formal analysis: A.S.P., I.A. S.E.D.; Intervention: W.A., O.H., A.P; PhosphoProteomic Investigation: H.H., L.H.H.L, T.C., R.J.M., S.J.H., I.A., A.S.P.; Proteomic Investigation: I.A, A.S.P. S.E.D.; Visualisations: A.S.P.; Funding acquisition: W.A., A.P., O.H.; Writing – original draft: I.A, A.P., A.S.P.; Writing – review and editing: All authors.

## Ethics declarations

The authors declare no competing interests.

**Extended Figure 1:**
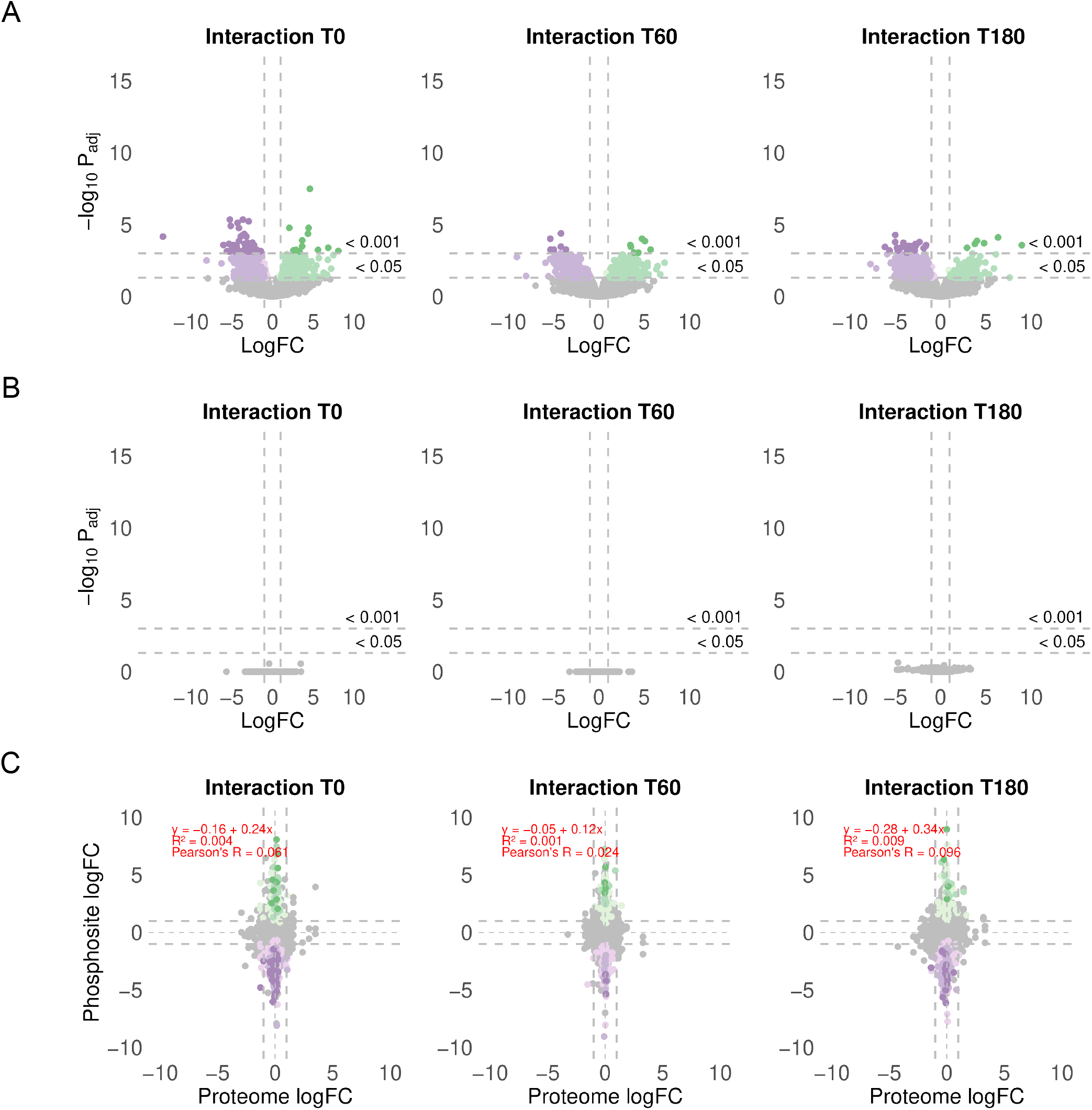
Interaction effects old vs young. **A**. Identified phosphosites from interactions between old and young at T0, T60 and T180. X-axis is the log fold change and Y-axis −log10 adjusted p-values. **B**. Identified proteins from interactions between old and young T0, T60 and T180 compared with baseline. X-axis is the log fold change and Y-axis −log10 adjusted p-values in young and old. **C**. Phosphosite logFC vs proteome logFC for T0, T60 and T180 interactions. Red dashed line is the linear regression fit.

**Extended Figure 2:**
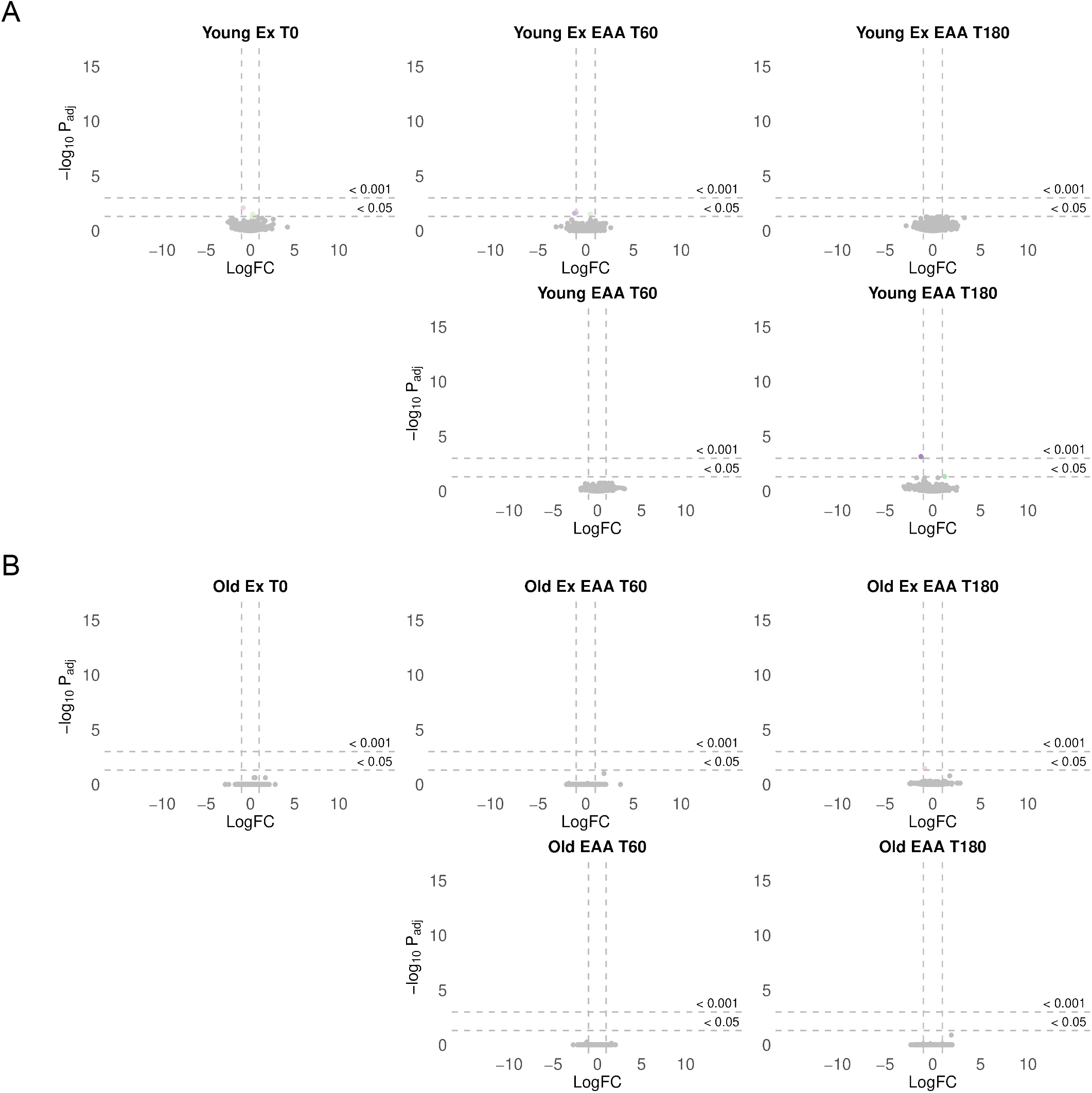
Proteome results for exercised conditions vs baseline in young and old participants. **A-B**Identified proteins from T0, T60 and T180 compared with baseline. X-axis is the log fold change and Y-axis −log10 adjusted p-values in young and old. Grey dashed lines on all plots indicate logFC |1| or adjusted P value cut off <0.05 and <0.001, and all blue points indicate downregulated and red upregulated.

**Extended Figure 3:**
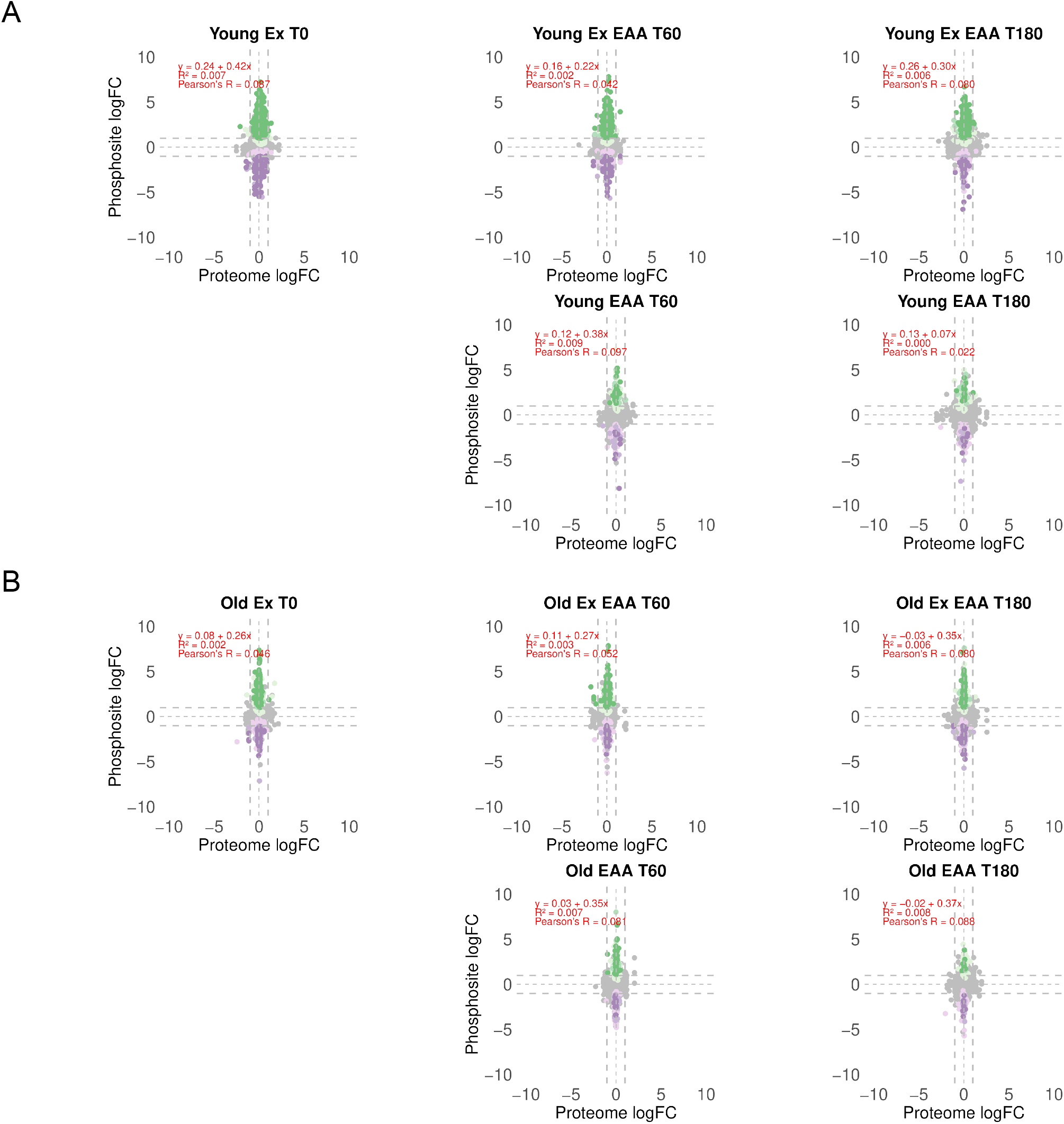
Correlation plots for exercised conditions vs baseline in young and old participants. **A-B** Phosphosite logFC vs proteome logFC for T0, T60 and T180. Red dashed line is the linear regression fit. Grey dashed lines on all plots indicate logFC |1|

**Extended Figure 4:**
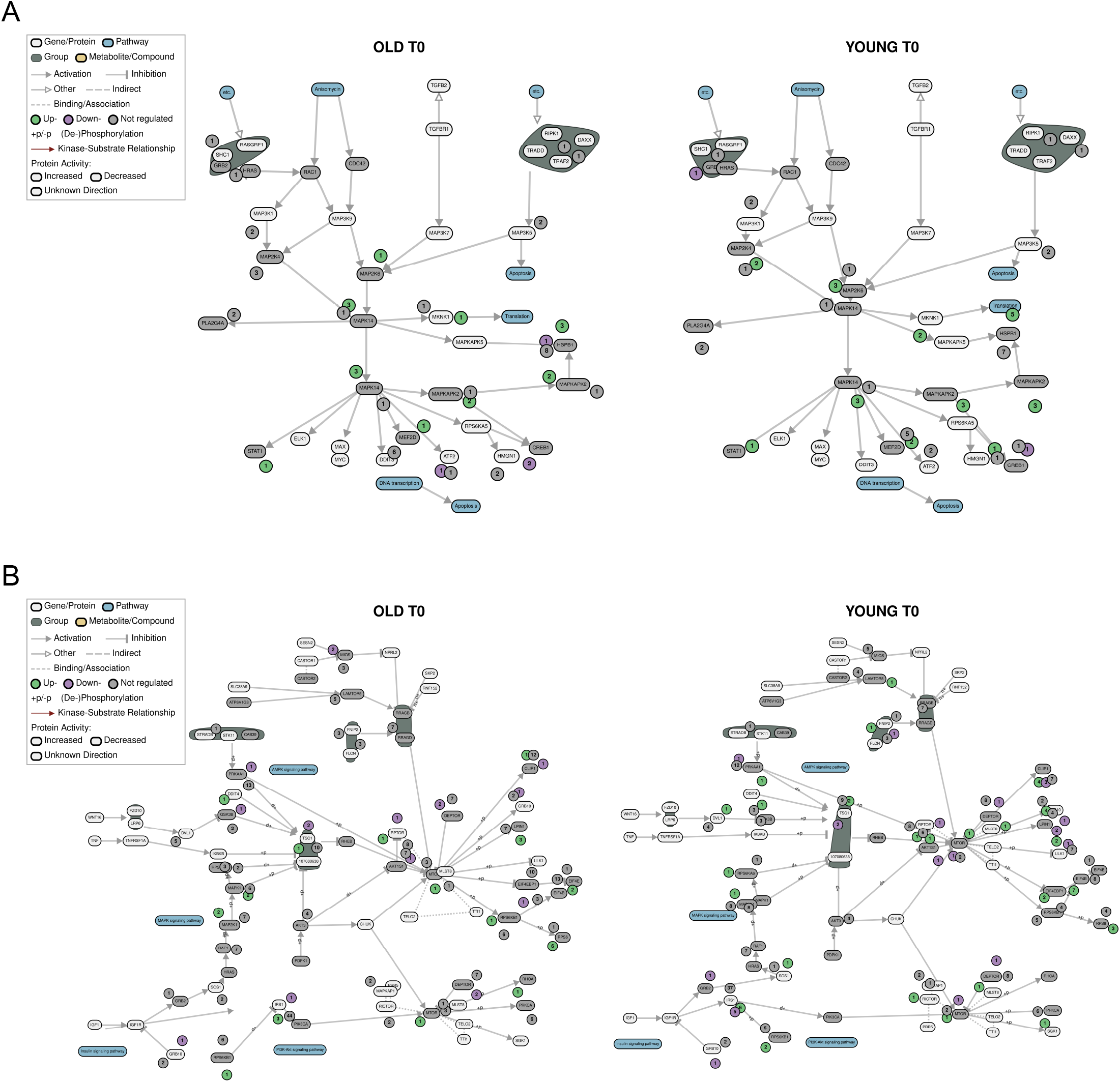
PTM Navigator mTOR signalling plot **A**. PTM navigator plot for the mTOR (HSA04150) signalling pathway (hsa04150). Phosphorylated sites indicated by small circle with numbers of sites, purple are down regulated, green upregulated and grey unregulated phosphosites. Protein regulation is shown by green upregulated, purple downregulated and grey unregulated. **B**. PTM navigator plot for the p38 MAPK (WP400) signalling pathway (hsa04150). Phosphorylated sites indicated by small circle with numbers of sites, purple are down regulated, green upregulated and grey unregulated phosphosites. Protein regulation is shown by green upregulated, purple downregulated and grey unregulated.

## Notes

https://doi.org/10.5281/zenodo.20387916

http://proteomecentral.proteomexchange.org

### Competing Interest Statement

The authors have declared no competing interest.

## References

1 Brook MS, Wilkinson DJ, Phillips BE, Perez-Schindler J., Philp A, Smith K et al. Skeletal muscle homeostasis and plasticity in youth and ageing: impact of nutrition and exercise. Acta physiologica 2015; 216: 15–41.

2 Anker SD, Morley JE, von Haehling S. Welcome to the icd-10 code for sarcopenia. Journal of cachexia, sarcopenia and muscle 2016; 7: 512–514.

3 Cruz-Jentoft AJ, Sayer AA. Sarcopenia. The lancet 2019; 393: 2636–2646.

4 Kumar V, Selby A, Rankin D, Patel R, Atherton P, Hildebrandt W et al. Age-related differences in the dose–response relationship of muscle protein synthesis to resistance exercise in young and old men. The journal of physiology 2009; 587: 211–217.

5 Cuthbertson D, Smith K, Babraj J, Leese G, Waddell T, Atherton P et al. Anabolic signaling deficits underlie amino acid resistance of wasting, aging muscle. The faseb journal 2004; 19: 1–22.

6 Morton RW, Traylor DA, Weijs PJ, Phillips SM. Defining anabolic resistance: implications for delivery of clinical care nutrition. Current opinion in critical care 2018; 24: 124–130.

7 Horwath O, Moberg M, Hodson N, Edman S, Johansson M, Andersson E et al. Anabolic sensitivity in healthy, lean, older men is associated with higher expression of amino acid sensors and mtorc1 activators compared to young. Journal of cachexia, sarcopenia and muscle 2024; 16. doi:10.1002/jcsm.13613.

8 Yilmaz S, Ayati M, Schlatzer D, Çiçek AE, Chance MR, Koyutürk M. Robust inference of kinase activity using functional networks. Nature communications 2021; 12. doi:10.1038/s41467-021-21211-6.

9 Humphrey SJ, Karayel O, James DE, Mann M. High-throughput and high-sensitivity phosphoproteomics with the easyphos platform. Nature protocols 2018; 13: 1897–1916.

10 Rappsilber J, Ishihama Y, Mann M. Stop and go extraction tips for matrix-assisted laser desorption/ionization, nanoelectrospray, and lc/ms sample pretreatment in proteomics. Analytical chemistry 2002; 75: 663–670.

11 R Core Team. R: A language and environment for statistical computing. R Foundation for Statistical Computing: Vienna, Austria, 2025 https://www.R-project.org/.

12 Bruderer R, Bernhardt OM, Gandhi T, Miladinović SM, Cheng L-Y, Messner S et al. Extending the limits of quantitative proteome profiling with data-independent acquisition and application to acetaminophen-treated three-dimensional liver microtissues. Molecular & cellular proteomics 2015; 14: 1400–1410.

13 Ritchie ME, Phipson B, Wu D, Hu Y, Law CW, Shi W et al. Limma powers differential expression analyses for rna-sequencing and microarray studies. Nucleic acids research 2015; 43: e47.

14 Smyth GK. Linear models and empirical bayes methods for assessing differential expression in microarray experiments. Statistical applications in genetics and molecular biology 2004; 3: 1–25.

15 Benjamini Y, Hochberg Y. Controlling the false discovery rate: A practical and powerful approach to multiple testing. Journal of the royal statistical society series b: Statistical methodology 1995; 57: 289–300.

16 Korotkevich G, Sukhov V, Budin N, Shpak B, Artyomov MN, Sergushichev A. Fast gene set enrichment analysis. 2016. doi:10.1101/060012.

17 Müller J, Bayer FP, Wilhelm M, Schuh MG, Kuster B, The M. Ptmnavigator: interactive visualization of differentially regulated post-translational modifications in cellular signaling pathways. Nature communications 2025; 16. doi:10.1038/s41467-024-55533-y.

18 Slowikowski K. Ggrepel: Automatically position non-overlapping text labels with “ggplot2”. Cran: Contributed packages. 2016. doi:10.32614/cran.package.ggrepel.

19 Gu Z, Eils R, Schlesner M. Complex heatmaps reveal patterns and correlations in multidimensional genomic data. Bioinformatics 2016; 32: 2847–2849.

20 Wilke CO. Cowplot: Streamlined plot theme and plot annotations for “ggplot2”. Cran: Contributed packages. 2015. doi:10.32614/cran.package.cowplot.

21 Graumann J, Cotton R. Multipanelfigure: Simple assembly of multiple plots and images into a compound figure. Journal of statistical software 2018; 84. doi:10.18637/jss.v084.c03.

22 Perez-Riverol Y, Bandla C, Kundu DJ, Kamatchinathan S, Bai J, Hewapathirana S et al. The pride database at 20 years: 2025 update. Nucleic acids research 2024; 53: D543–D553.

23 Palmer AS, Alldritt I, Apro W, Huckstep H, Le L, Horwath O et al. Ageing blunts the phospho-proteomic response to resistance exercise in humans - data and code. 2026. doi:10.5281/zenodo.20387916.

